# Juvenile emperor penguin range calls for extended conservation measures in the Southern Ocean

**DOI:** 10.1101/2021.04.06.438390

**Authors:** Aymeric Houstin, Daniel P. Zitterbart, Karine Heerah, Olaf Eisen, Víctor Planas-Bielsa, Ben Fabry, Céline Le Bohec

## Abstract

To protect the unique Southern Ocean biodiversity, conservation measures like marine protected areas (MPAs) are implemented based on the known habitat distribution of ecologically important species. However, distribution models focus on adults, neglecting that immatures animals can inhabit vastly different areas. Here, we show that current conservation efforts in the Southern Ocean are insufficient for ensuring the protection of the highly mobile Emperor penguin. We find that juveniles spend ∼90% of their time outside the boundaries of proposed and existing MPAs, and that their distribution extends far beyond (> 1500 km) the species’ extent of occurrence as defined by the International Union for Conservation of Nature. We argue that strategic conservation plans for Emperor penguin and long-lived ecologically important species must consider the dynamic habitat range of all age classes.

## Introduction

Anthropogenic environmental changes lead to upheaval even in remote and apparently untouched ecosystems such as the Antarctic and the Southern Ocean. Marine top predators, such as seabirds and marine mammals, play a pivotal role in marine ecosystems ^1^, and any disruptions in their abundance and distribution can have a major impact on the functioning and resilience of ecosystems ^2^. At the same time, top predators are indicators of ecosystem health because of their high position in the trophic cascade and the vast, ocean basin-scale habitat of individual animals ^3,4^. Thus, top predators integrate signals from across the food web and are therefore important bioindicators ^5^. The health, abundance and distribution of marine top predators are consequently key metrics in ecosystem-based management and systematic conservation planning ^6^.

Effective conservation plans require comprehensive consideration of the at-sea distribution of species, including each life-history stage such as juveniles and immatures as they constitute an essential part of the total population ^7^. However, in some of these taxa, in particular in many seabird species, the distribution of juveniles and immatures is difficult to assess and is therefore often neglected. This is especially true for polar ecosystems, where remoteness and the extreme environmental conditions hamper data collection.

Currently, the Southern Ocean experiences significant impacts due to global change ^8,9^. Measurable negative effects on wildlife have already occurred, such as population decreases of numerous seabird species ^10,11^, including the complete loss of emperor penguin (*Aptenodytes forsteri)* colonies ^12,13^. The vanishing of these colonies has been attributed to strong El Niño events, rise in local mean annual air temperature, strong winds, and/or decline in seasonal sea ice duration. Climate change is also expected to result in human access to new ice-free fishing areas ^14^, whereby seabirds and marine mammals will have to compete for food with industrial fisheries and may even become by-catch ^15^. The accumulation of anthropogenic pressures on these fragile ecosystems urgently requires effective protection ^16^.

The Commission for the Conservation of Antarctic Marine Living Resources (CCAMLR) is the governing body in charge of conservation issues in the Southern Ocean. CCAMLR’s mandate includes the implementation of conservation measures, such as the establishment of Marine Protected Areas (MPAs) and the regulation of the fishing industry, through quota allocations and gear limitations ^17^. Within the CCAMLR, conservation measures are based on the best scientific data available, including the distribution and demography of marine predators ^18,19^. Similarly, the International Union for Conservation of Nature (IUCN)’s Red List of Threatened Species depicts the extent of occurrence (EOO) of each species, i.e. all the known, inferred or projected sites of present occurrence of the species’ adults excluding cases of vagrancy ^20^. Such knowledge serves then as a reference for policy making on the implementation of conservation measures. Consequently, providing novel data, in areas that have never been surveyed or on data-deficient population classes like juveniles enhances the conservation governance perspective for a species and its habitat.

Currently, 12% of the waters inside the CCAMLR boundaries are protected, with only 4.6% as no-take areas. This includes the Ross Sea and waters around South Orkney Islands ^16^. Since 2002, the CCAMLR has been working on establishing a network of MPAs around Antarctica, but the implementation of three new MPAs in East Antarctica, the Weddell Sea, and at the Antarctic Peninsula has been difficult. But even when implemented, the new MPAs would protect only 22% of the Southern Ocean inside the CCAMLR boundaries^16^, which is significantly less than the IUCN recommended protection target of 30% of each marine habitat ^21^. Furthermore, assessments and recommendations are based on limited and incomplete data. For instance, in the Weddell Sea, home to one-third of the global emperor penguin population, no tracking studies have been conducted so far; thus, very little is known about the penguins’ at-sea distribution in this area.

The Emperor penguin is considered an iconic and ecologically important species of Antarctica. Its colony sites and at-sea movements have been the basis of previous discussions of conservation priorities, either in terms of MPAs ^4^, Important Bird Areas ^22,23^ or Areas of Ecological Significance (AESs ^24^). With a population currently estimated at *ca*. 270 000 breeding pairs in 61 known colonies around the continent ^25^, the species is severely threatened by global warming and expanding fishing activities in the Southern Ocean ^15,26^, facing the risk to be nearly extinct within this century ^27^. The most effective actions to protect the Emperor penguin from anthropogenic impacts would be a reduction in greenhouse gas emissions ^26,27^ as well as the establishment of MPAs throughout its habitat range ^26^. However, little is known about the early life at sea of emperor penguins, even though their survival is crucial for the viability of the global population ^28^. To date, a total of only 48 juvenile emperor penguins have been tracked. Moreover, tracking has been done only in the Ross Sea and East Antarctica (Table 1), even though for the designation of MPAs, it is fundamental to know their distribution at the circum-Antarctic scale ^4,6,7^.

**Table 1.**
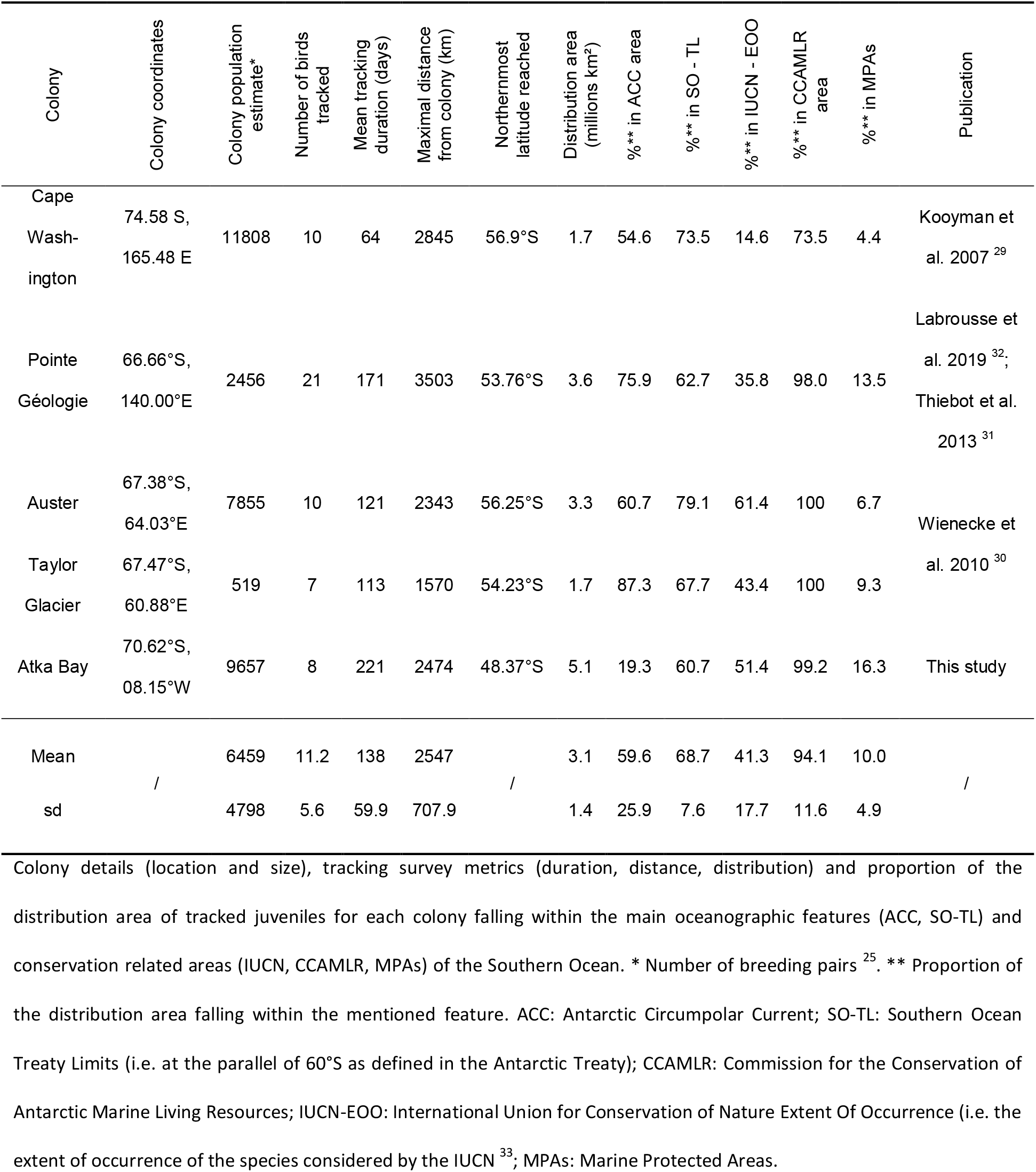
Tracking studies of juvenile emperor penguins at sea.

The aim of this study was to bridge this gap in knowledge by equipping 6-months-old emperor penguin chicks with ARGOS satellite platforms that transmit the birds’ locations several times each day. Birds were tagged before their initial departure from their colony of origin at Atka Bay (70°37’S, 08°09’W) near the south eastern limit of the Weddell Sea (Fig. 1). We recorded their journey during their first year at sea (Fig. 1 and Supplementary Table 1). To assess the habitat range used by the juvenile emperor penguins at the scale of the Southern Ocean, we incorporated the distribution of all previously tracked juvenile emperor penguins into our analysis (Fig. 1; ^29–32^). We find that juveniles travel beyond the boundaries of existing and planned conservation and management areas, demonstrating that conservation efforts in the Southern Ocean are insufficient to protect emperor penguins.

**Fig. 1.**
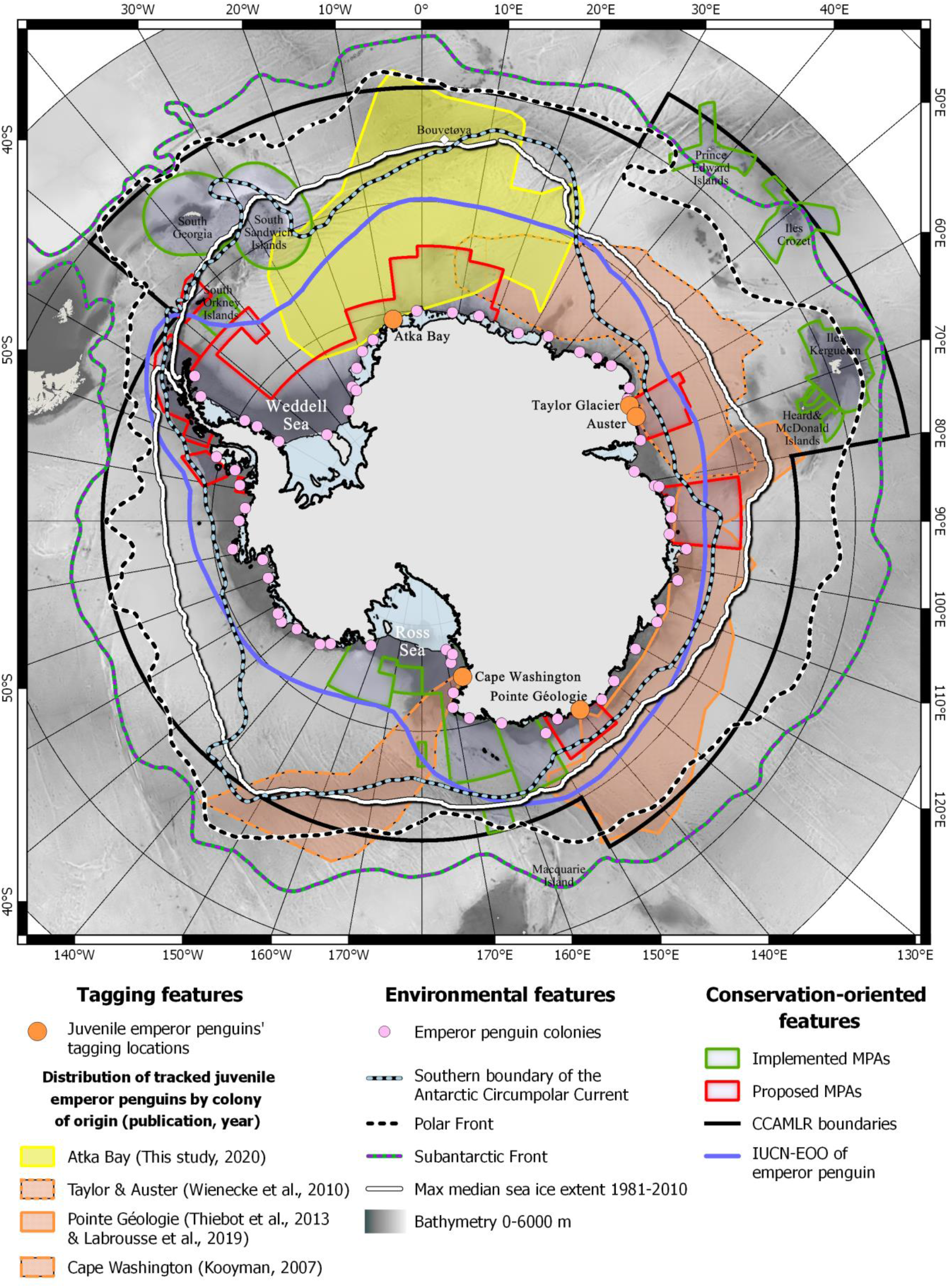
Overlap between existing and planned conservation zones and the distribution of juvenile emperor penguins tracked to date in the Southern Ocean. Distribution areas of juveniles are indicated by colored polygons. MPAs: Marine Protected Areas; CCAMLR: Commission for the Conservation of Antarctic Marine Living Resources; IUCN-EOO: International Union for Conservation of Nature Extent Of Occurrence (i.e. the extent of occurrence of the species considered by the IUCN ^33^).

## Results

The tracking data from our study show that juvenile emperor penguins travelled north of 50°S (the lowest recorded latitude was 48.37°S), which is 600 km further north than previously recorded (Table 1). Two of the eight tagged birds reached the South Sandwich Islands region in winter (late June until at least July) before their ARGOS platforms stopped transmitting. Thus, with three of the eight birds reaching sub-Antarctic areas, the presence of juvenile emperor penguins in these waters should be considered common rather than unusual behavior. All tagged juveniles reached the southern boundary of the Antarctic Circumpolar Current (ACC), and five of the tagged birds remained between the southern ACC boundary and the Antarctic Polar Front for prolonged time periods (> 46 days). One bird travelled north of the Polar Front ^29,32^. The penguin tracks over a full year (polygon encompassing the area covered by the tracks, Supplementary Fig. 1) covered an area of 5.1 million km2 (Fig. 1 and Fig. 2, Table 1), nearly 1.4 times larger than the largest previously reported distribution of juvenile emperor penguins from their colony of origin (Table 1).

**Fig. 2.**
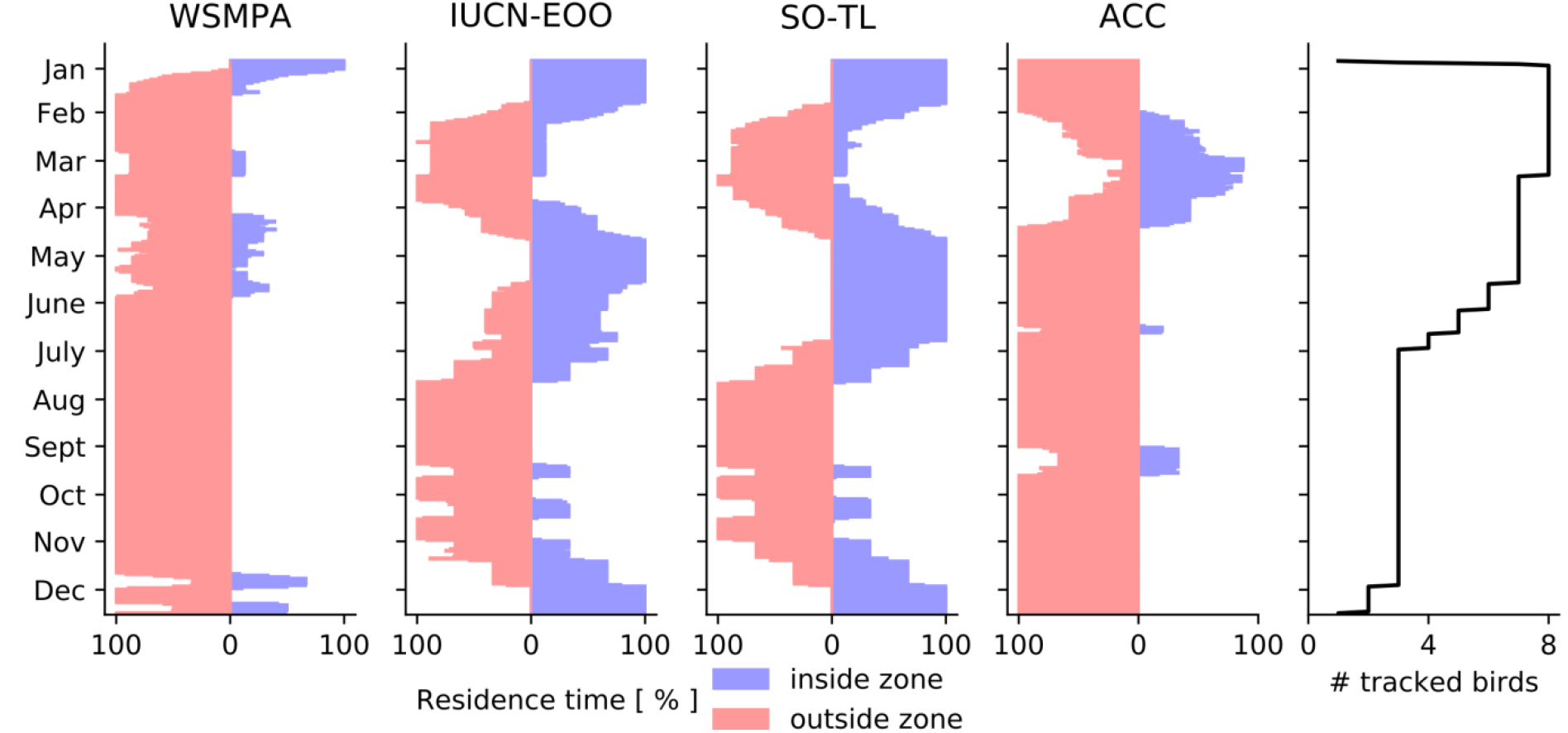
Proportion of time that the eight tagged juvenile emperor penguins from the Atka Bay colony spent either inside or outside the main conservation related areas (WSMPA, IUCN-EOO) and oceanographic features (SO-TL, ACC) of the Atlantic sector of the Southern Ocean. Daily average across all individuals computed over hourly data points. WSMPA: Weddell Sea Marine Protected Area; IUCN-EOO: International Union for Conservation of Nature Extent Of Occurrence (i.e. the extent of occurrence of the species considered by the IUCN ^33^; SO-TL: Southern Ocean Treaty Limits (i.e. at the parallel of 60°S as defined in the Antarctic Treaty); ACC: Antarctic Circumpolar Current.

Juvenile emperor penguins from the Weddell Sea area had a seasonal travel pattern similar to that of those tracked in other sectors of the Southern Ocean. After leaving their colony, juveniles migrated northward towards and into the ACC where they remained for 37 ± 24 days. Juvenile emperor penguins commonly ranged outside the limits of the Southern Ocean (i.e. the parallel of 60°S as defined by the Antarctic Treaty, hereafter referred to as SO-Treaty); some birds travelled outside the CCAMLR boundaries (Fig. 1, Supplementary Table 1). Over the course of April, juveniles migrated southward towards the pack ice to spend the winter (Supplementary Fig. 2). In this study, the juvenile penguins spent only 51.1 ± 13.3% of their time inside the IUCN-EOO of the species, which is based on the estimated adult distribution (Supplementary Fig. 2). Moreover, the time spent inside the area varied significantly across months (p<1e-05; Fig. 2). In August (winter), all penguins were outside and travelled up to 1260 km north of the IUCN-EOO, whereas they were mostly inside in January and May. When considering the data from all studies, 41.3 ± 17.7% of the observed distribution areas of juvenile emperor penguins fell within the IUCN-EOO (Table 1). Juveniles from the Cape Washington colony in the Ross Sea travelled up to 1500 km outside the IUCN-EOO.

Taken together, the EOO of emperor penguin defined by the IUCN is underestimating the current habitat range of the species. Existing and planned MPAs cover on average only 10.0 ± 4.9% of the estimated distribution areas (Table 1). Regarding the time spent inside protected areas, juvenile emperor penguins from the Atka Bay colony, which is located inside the proposed Weddell Sea Marine Protected Areas (WSMPA, the largest currently proposed MPA in the Southern Ocean), left the MPA’s boundaries after 9 ± 4 days in January, and remained only 10.6 ± 7.5% of their time inside the boundaries (Fig. 2). Only during summer (January and December) did the juveniles spend a considerable amount of time inside the WSMPA (47.9 ± 23.8% and 31.1 ± 13.4%, respectively). All tagged penguins were outside the WSMPA’s boundaries in February and from July to November (Fig. 2).

## Discussion

Penguins are considered umbrella species of the Southern Ocean’s ecosystem ^34^. Monitoring their population trend and distribution is therefore essential for biodiversity conservation. The common approach for designating boundaries of MPAs focuses on protecting the breeding segment of populations ^35^. We argue that this might not be sufficient for species for which juvenile and adult ranges do not overlap, and we point out that the habitat range of juvenile penguins requires also a high level of protection. Indeed, juveniles are more vulnerable than adults as their foraging skills (including their ability to dive, to capture prey, and to find productive feeding grounds) are not yet fully developed, and their experience to escape predators is minimal ^36^. Moreover, juvenile survival can have a critical impact on the population dynamics, especially in long-lived species ^37^. Emperor penguins start breeding earliest at age 4-5 years, lay only one egg per pair and year, and only have an annual chance of 55% to bring a chick to fledging ^27^. This low fecundity, projected to decrease under future warming scenarios ^27^, makes the survival of immature individuals, which represent about one quarter of the total population ^28^, particularly critical for the recruitment into breeding populations and thus the species’ viability ^38^. Moreover, in contrast to adults, the dispersal behavior of juveniles is one of the main processes by which long-lived species will be able to adapt to the ongoing rapid environmental change. A vast travel range allows them to explore possible alternative feeding and breeding grounds ^39^. Therefore, for successful conservation we need to consider the habitat range of all age classes.

Our findings reveal that juveniles commonly spent a considerable amount of time outside the species’ IUCN-EOO and outside the limits of existing or planned MPAs in the Southern Ocean (Fig. 1, Table 1 and Supplementary Table 1). Consequently, if protection measures were based solely on the current IUCN-EOO of the species, as it stands, given its focus on adult occurrences due to the insufficient data for juveniles, this could lead to inefficient decisions for the future protection of the species. Furthermore, all studies including ours have reported that juveniles visit the highly productive ACC area during their first journey at sea, where the Antarctic Polar Front appears to act as an ecological barrier. During the most vulnerable stage of their life, the penguins’ dispersive behavior leads them outside the SO-Treaty and CCAMLR limits into waters where they are likely to encounter and compete with fisheries (see ^23,40^ for data on fisheries activity). In accordance with the CCAMLR’s ecosystem-based fisheries management approach, the presence of this critical fragment of the emperor penguin population should be considered by the CCAMLR when allocating fishing quotas and zones; especially in the current context where several CCAMLR fishing states are lobbying for an increase of the spatial and temporal distribution of catches and fisheries ^41^.

A growing body of evidence indicates the ongoing threats to penguins. Trathan and colleagues ^26^ recently advocated for a reclassification of the Emperor penguin on the IUCN Red List from the current “Near Threatened” status to “Vulnerable” or “Endangered”, together with the classification as an “Antarctic Specially Protected Species” by the Antarctic Treaty. Our data support this call for better protection by also pointing out the need to include all age-classes and age-specific threats into the classification assessment ^4,7^. Furthermore, in the context of the vast range of emperor penguins and other marine top predators ^4^, our data argue in favor of a circumpolar integrated systems of marine protected areas in the Southern Ocean that consider large parts of the offshore Antarctic Convergence area. This could be achieved, for instance by combining migratory corridors with static and dynamic MPAs (i.e. MPAs that rapidly evolve in space and time in response to changes in the ocean and its users; ^42^), to create an ecological connected network ^43^ that would provide more effective protection to the Southern Ocean ecosystem ^16^.

## Materials and Methods

### Study site and instrumentation

Our study was conducted at the Atka Bay emperor penguin colony (70°37’S, 08°09’W) near (∼ 10 km) the German Antarctic research base “Neumayer Station III”. In January 2019, we equipped eight 6-month-old chick emperor penguins with satellite communicating SPOT-367 ARGOS platforms (Wildlife Computers, Redmond, WA 98052, USA). The ARGOS platforms were programmed to transmit their identification every day at 4, 6, 10, 16, 19 and 21:00 GMT, corresponding to time points with optimum ARGOS satellite coverage over the Weddell Sea (ARGOS CLS, Toulouse, France).

To minimize drag, the ARGOS platforms were deployed on the lower back of the birds ^44,45^. The streamlined devices were attached to the feather with adhesive tape (Tesa tape 4651, Beiersdorf AG, Hamburg, Germany) and secured with three cable ties (Panduit PLTM1.5M-C0 142*2.6 mm, Panduit Corp, Illinois, USA). We then applied epoxy glue (Loctite EA 3430, Loctite, Henkel AG., Düsseldorf, Germany) on the mounting to increase waterproofing and robustness ^46,47^.

### Estimation of the at-sea distribution of juvenile emperor penguins from the Atka Bay colony

#### Location filtering

ARGOS locations are associated with spatial error ellipses. These spatial errors can range from a few hundred meters to several kilometers ^48,49^. Erroneous locations were filtered out using a speed filter from the R package ‘*argos filter*’ ^50^ with the maximum travel speed fixed at 15 km/h following similar studies on emperor penguins ^32,51^.

#### Interpolation of locations at a regular time step

We used a state-space modelling approach ^52^ to estimate hourly locations. Specifically, a Kalman filter, which accounted for location error, was applied using the R package ‘*crawl*’ ^53^, and Continuous-time Correlated Random Walk (CRW) models were used to predict locations at a regular time step interval of 1 h ^52,54^.

### Estimation of the colony-specific distribution area for juvenile emperor penguins

In addition to the 8 birds tracked in our study, 48 juvenile emperor penguins from 4 different colonies were previously tracked ^29–32^, see Table 1 for the details on the colonies). Data of these previously acquired bird journeys are available as maps in the respective publications. We georeferenced these tracking maps using the QGIS software. We subsequently plotted the main corner points encompassing the tracks of all birds from each colony (Supplementary Fig. 1). We obtained the distribution of juvenile emperor penguins by computing the concave hull envelope for each dataset using the ‘*ConcaveHull’* plugin ^55^. Envelopes from the same colony ^31,32^ were merged to consider only one polygon per colony (referred to as distribution area), including one for the Atka Bay colony. The size of each distribution area was calculated with the ‘*raster’* package in R ^56^ and is reported in Table 1. Due to the significant overlap of Auster and Taylor Glacier juvenile distribution ^30^ and the proximity (132 km) of the two sites ^57^, for visualization purposes, the tracks of the birds from Auster and Taylor Glacier colonies are shown in the same polygon in Fig. 1. However, the distribution areas were computed separately for each colony.

### Ecological features

The locations of the Southern Ocean fronts and the Antarctic Circumpolar Current boundaries (ACC, ^58^) were downloaded from https://gis.ccamlr.org ^59^.

The bathymetry at one-minute horizontal spatial resolution was obtained from the ETOPO1 Global Relief Model provided by the NOAA National Geophysical Data Center ^60^.

Sea ice concentrations (ranging from 0-100%) were obtained from Advanced Microwave Scanning Radiometer (AMSR-2) satellite estimates of daily sea ice concentration at 3.125 km resolution from the University of Bremen (https://seaice.uni-bremen.de/data/amsr2/) ^61^. The sea ice edge contour was defined by the 15% sea ice concentration ^62,63^ (Supplementary Fig. 2).

The maximum and minimum median sea ice extent from 1981-2010 presented in Supplementary Fig. 1 and Supplementary Fig. 2 were obtained from the National Snow and Ice Data Center NSDIC ^64^ implemented in the ‘*Quantarctica3’* package ^65^ of the QGIS software.

### Conservation oriented features

The Commission for the Conservation of Antarctic Marine Living Resources (CCAMLR) planning domains and existing Antarctic Marine Protected Areas (MPAs) were obtained from https://gis.ccamlr.org ^59^. The proposed Weddell Sea Marine Protected Area boundaries (WSMPA, ^66^) and the proposed East Antarctic Marine Protected Area boundaries (EAMPA, ^67^) were obtained from www.mpatlas.org ^68^. The Domain 1 MPA proposal ^69^ was drawn from www.mpatlas.org ^68^. The South Georgia and South Sandwich Islands Marine Protected Area (SGSSIMPA) and the sub-Antarctic MPAs boundaries were downloaded from www.protectedplanet.net.

The International Union for Conservation of Nature (IUCN) extent of occurrence (EOO) of the Emperor penguin species was obtained from www.iucnredlist.org ^70^.

### Assessing the overlap between bird distribution and conservation oriented areas

The average residency time that each of the birds equipped in our study spent inside existing or proposed conservation oriented areas of the Southern Ocean was computed on a daily, weekly or monthly basis, or averaged over the total tracking period.

We tested whether the observed monthly-averaged residency time changed significantly over the course of a year using the Kruskal-Wallis rank sum tests. For all tests, the significance threshold was set at p=0.05. Statistical analyses were performed using the software R v. 3.5.0 ^71^ and QGIS v. 2.18.18 ^72^ with the data package ‘*Quantarctica3*’ ^65^.

## Data and materials availability

All data generated or analysed during this study will be available in the Movebank data repository at https://www.movebank.org/.

## Acknowledgements

We thank the Alfred-Wegener-Institut Helmholtz-Zentrum für Polar-und Meeresforschung (AWI), Logistics Department, the winterers and campaigners at Neumayer Station III for their invaluable support. We are very grateful to Prof. Patrick Rampal for support in initiating the project. This study was funded by the Centre Scientifique de Monaco with additional support from the LIA-647 and RTPI-NUTRESS (CSM/CNRS-University of Strasbourg), by The Penzance Endowed Fund and The Grayce B. Kerr Fund in Support of Assistant Scientists, and by the Deutsche Forschungsgemeinschaft (DFG) grants ZI1525/3-1 in the framework of the priority program “Antarctic research with comparative investigations in Arctic ice areas”. Logistics and field efforts were supported by the AWI within the framework of the program “Monitor the health of the Antarctic maRine ecosystems using the Emperor penguin as a sentinel” (MARE). The long-term project MARE, to which this study belongs, and all procedures were approved by the German Environment Agency (Umweltbundesamt-UBA permit no.: II 2.8 – 94033/100 delivered on the 04/10/2017 and 04/10/2018), and conducted in accordance with the Committee for Environmental Protection (CEP) guidelines.

## Funding

This study was funded by the Centre Scientifique de Monaco with additional support from the LIA-647 and RTPI-NUTRESS (CSM/CNRS-University of Strasbourg), by The Penzance Endowed Fund and The Grayce B. Kerr Fund in Support of Assistant Scientists, and by the Deutsche Forschungsgemeinschaft (DFG) grants ZI1525/3-1 in the framework of the priority program “Antarctic research with comparative investigations in Arctic ice areas”.

## Competing interests

Authors declare no competing interests.

## Author contributions

AH and CLB conceived the ideas and designed the methodology and protocols. AH, OE and CLB, conducted fieldwork. AH and KH analysed data. AH, DZ, OE and CLB interpreted the results. AH, DZ, BF and CLB led the writing of manuscript. All authors edited and proofread the paper.

## Notes

https://www.movebank.org

### Competing Interest Statement

The authors have declared no competing interest.

## References

1. Pimiento, C. et al. Functional diversity of marine megafauna in the Anthropocene. Science Advances 6, eaay7650 (2020).

2. Baum, J. K. & Worm, B. Cascading top-down effects of changing oceanic predator abundances. Journal of Animal Ecology 78, 699–714 (2009).

3. Hazen, E. L. et al. Marine top predators as climate and ecosystem sentinels. Frontiers in Ecology and the Environment 17, 565–574 (2019).

4. Hindell, M. A. et al. Tracking of marine predators to protect Southern Ocean ecosystems. Nature 580, 87–92 (2020).

5. Trathan, P. N., Forcada, J. & Murphy, E. J. Environmental forcing and Southern Ocean marine predator populations: effects of climate change and variability. Philosophical Transactions of the Royal Society B: Biological Sciences 362, 2351–2365 (2007).

6. Hays, G. C. et al. Translating marine animal tracking data into conservation policy and management. Trends in Ecology & Evolution 34, 459–473 (2019).

7. Carneiro, A. P. B. et al. A framework for mapping the distribution of seabirds by integrating tracking, demography and phenology. Journal of Applied Ecology 57, 514–525 (2020).

8. Stark, J. S., Raymond, T., Deppeler, S. L. & Morrison, A. K. Antarctic Seas. in World Seas: an Environmental Evaluation 1–44 (Elsevier, 2019).

9. Swart, N. C., Gille, S. T., Fyfe, J. C. & Gillett, N. P. Recent Southern Ocean warming and freshening driven by greenhouse gas emissions and ozone depletion. Nature Geoscience 11, 836–841 (2018).

10. Trivelpiece, W. Z. et al. Variability in krill biomass links harvesting and climate warming to penguin population changes in Antarctica. Proceedings of the National Academy of Sciences 108, 7625–7628 (2011).

11. Ropert-Coudert, Y. et al. Two recent massive breeding failures in an Adélie penguin colony call for the creation of a Marine Protected Area in D’Urville Sea/Mertz. Frontiers in Marine Science 5, 1–7 (2018).

12. Trathan, P. N., Fretwell, P. T. & Stonehouse, B. First recorded loss of an emperor penguin colony in the recent period of Antarctic regional warming: implications for other colonies. PLoS ONE 6, e14738 (2011).

13. Fretwell, P. T. & Trathan, P. N. Emperors on thin ice: three years of breeding failure at Halley Bay. Antarctic Science 31, 133–138 (2019).

14. Rintoul, S. R. et al. Choosing the future of Antarctica. Nature 558, 233–241 (2018).

15. Trathan, P. N. et al. Pollution, habitat loss, fishing, and climate change as critical threats to penguins. Conservation Biology 29, 31–41 (2015).

16. Brooks, C. M. et al. Progress towards a representative network of Southern Ocean protected areas. PLOS ONE 15, e0231361 (2020).

17. CCAMLR. Schedule of conservation measures in force 2020/21. (2020).

18. CCAMLR. Resolution 31/XXVIII, Best Available Science. 1–2 (2009).

19. Teschke, K. et al. An integrated data compilation for the development of a marine protected area in the Weddell Sea. Earth System Science Data Discussions 1–31 (2019).

20. IUCN. IUCN Red List Categories and Criteria: Version 3.1. (2012).

21. IUCN. The Promise of Sydney, IUCN World Parks Congress. Sydney, Australia. (2014).

22. Harris, C.M., Lorenz, K., Fishpool, L.D.C., Lascelles, B., Cooper, J., Coria, N.R., Croxall, J. P. & Emmerson, L.M., Fraser, W.R., Fijn, R.C., Jouventin, P., LaRue, M.A., Le Maho, Y., Lynch, H.J., Naveen, R., Patterson-Fraser, D.L., Peter, H.-U., Poncet, S., Phillips, R.A., Southwell, C.J., van Franeker, J.A., Weimerskirch, H., Wienecke, B., & Woehler, E. J. Important Bird Areas in Antarctica 2015. BirdLife International and Environmental Research & Assessment Ltd., Cambridge (2015).

23. Handley, J. et al. Marine Important Bird and Biodiversity Areas for Penguins in Antarctica, Targets for Conservation Action. Frontiers in Marine Science 7, (2021).

24. Hindell, M. A. et al./person-group>. Foraging habitats of top predators, and Areas of Ecological Significance, on the Kerguelen Plateau. in The Kerguelen Plateau: marine ecosystem and fisheries (eds. Duhamel, G. & Welsford, D.) 35, 203–215 (2011).

25. Fretwell, P. T. & Trathan, P. N. Discovery of new colonies by Sentinel2 reveals good and bad news for emperor penguins. Remote Sensing in Ecology and Conservation rse2.176 (2020).

26. Trathan, P. N. et al. The emperor penguin - Vulnerable to projected rates of warming and sea ice loss. Biological Conservation 241, 108216 (2020).

27. Jenouvrier, S. et al. The Paris Agreement objectives will likely halt future declines of emperor penguins. Global Change Biology 26, 1170–1184 (2019).

28. Jenouvrier, S., Barbraud, C. & Weimerskirch, H. Long-term contrasted responses to climate of two antarctic seabird species. Ecology 86, 2889–2903 (2005).

29. Kooyman, G. L. & Ponganis, P. J. The initial journey of juvenile emperor penguins. Aquatic Conservation: Marine and Freshwater Ecosystems 17, S37–S43 (2007).

30. Wienecke, B., Raymond, B. & Robertson, G. Maiden journey of fledgling emperor penguins from the Mawson Coast, East Antarctica. Marine Ecology Progress Series 410, 269–282 (2010).

31. Thiebot, J.-B., Lescroël, A., Barbraud, C. & Bost, C.-A. Three-dimensional use of marine habitats by juvenile emperor penguins Aptenodytes forsteri during post-natal dispersal. Antarctic Science 25, 536–544 (2013).

32. Labrousse, S. et al. First odyssey beneath the sea ice of juvenile emperor penguins in East Antarctica. Marine Ecology Progress Series 609, 1–16 (2019).

33. IUCN. The IUCN Red List of Threatened Species. Version 2020-1. (2020). Available at: www.iucnredlist.org.

34. Le Bohec, C., Whittington, J. D. & Le Maho, Y. Polar monitoring: Seabirds as sentinels of marine ecosystems. in Adaptation and Evolution in Marine Environments From Pole to Pole (eds. Verde, C. & di Prisco, G.) 2, 205–230 (Springer Berlin Heidelberg, 2013).

35. Boersma, P. D. et al. Applying science to pressing conservation needs for penguins. Conservation Biology 34, 103–112 (2019).

36. Orgeret, F., Weimerskirch, H. & Bost, C.-A. Early diving behaviour in juvenile penguins: improvement or selection processes. Biology Letters 12, 20160490 (2016).

37. Stearns, S. C. The Evolution of Life Histories. (Oxford University Press, London, 1992).

38. Abadi, F., Barbraud, C. & Gimenez, O. Integrated population modeling reveals the impact of climate on the survival of juvenile emperor penguins. Global Change Biology 23, 1353–1359 (2017).

39. Gienapp, P. & Merilä, J. Evolutionary Responses to Climate Change. Encyclopedia of the Anthropocene 2, 51–59 (2018).

40. Brooks, C. M., Crowder, L. B., Österblom, H. & Strong, A. L. Reaching consensus for conserving the global commons: The case of the Ross Sea, Antarctica. Conservation Letters 13, 1–10 (2020).

41. Brooks, C. M. et al. Science-based management in decline in the Southern Ocean. Science 354, 185–187 (2016).

42. Maxwell, S. M. et al. Dynamic ocean management: Defining and conceptualizing real-time management of the ocean. Marine Policy 58, 42–50 (2015).

43. Allan, J. C., Beazley, K. F. & Metaxas, A. Ecological criteria for designing effective MPA networks for large migratory pelagics: Assessing the consistency between IUCN best practices and scholarly literature. Marine Policy 127, 104219 (2021).

44. Culik, B. & Wilson, R. P. Energetics of under-water swimming in Adélie penguins (Pygoscelis adeliae). Journal of Comparative Physiology B 161, 285–291 (1991).

45. Bannasch, R., Wilson, R. P. & Culik, B. Hydrodynamic aspects of design and attachment of a back-mounted device in penguins. The Journal of Experimental Biology 194, 83–96 (1994).

46. Wilson, R. P. et al. Long-term attachment of transmitting and recording devices to penguins and other seabirds. Wildlife Society Bulletin 25, 101–106 (1997).

47. Pütz, K. et al. Post-fledging dispersal of king penguins (Aptenodytes patagonicus) from two breeding sites in the South Atlantic. PLoS ONE 9, e97164 (2014).

48. Costa, D. P. et al. Accuracy of ARGOS locations of pinnipeds at-sea estimated using Fastloc GPS. PLoS ONE 5, e8677 (2010).

49. CLS. Argos User’s manual. (2016).

50. Freitas, C., Lydersen, C., Fedak, M. A. & Kovacs, K. M. A simple new algorithm to filter marine mammal Argos locations. Marine Mammal Science 24, 315–325 (2008).

51. Wienecke, B. & Robertson, G. Foraging space of emperor penguins Aptenodytes forsteri in Antarctic shelf waters in winter. Marine Ecology Progress Series 159, 249–263 (1997).

52. Johnson, D. S., London, J. M., Lea, M.-A. & Durban, J. W. Continuous-time correlated random walk model for animal telemetry data. Ecology 89, 1208–1215 (2008).

53. Johnson, D. S. Fit continuous-time correlated random walk models to animal movement data. 24 (2014).

54. Heerah, K. et al. Important areas and conservation sites for a community of globally threatened marine predators of the Southern Indian Ocean. Biological Conservation 234, 192–201 (2019).

55. Moreira, A. & Santos, M. Y. Concave hull: a k-nearest neighbours approach for the computation of the region occupied by a set of points. in Proceedings of the Second International Conference on Computer Graphics Theory and Applications GM, 61–68 (SciTePress - Science and and Technology Publications, 2007).

56. Hijmans, R. J. & van Etten, J. raster: Geographic analysis and modeling with raster data.?: R package version 3.1-5 (2020).

57. Ancel, A. et al. Looking for new emperor penguin colonies? Filling the gaps. Global Ecology and Conservation 9, 171–179 (2017).

58. Orsi, A. H., Whitworth, T. & Nowlin, W. D. On the meridional extent and fronts of the Antarctic Circumpolar Current. Deep-Sea Research Part I 42, 641–673 (1995).

59. CCAMLR. CCAMLR GIS 2019. (2019). Available at: https://gis.ccamlr.org.

60. Amante, C. & Eakins, B. W. ETOPO1 1 Arc-Minute Global Relief Model: Procedures, Data Sources and Analysis. NOAA Technical Memorandum NESDIS NGDC-24 19 (2009). doi:10.1594/PANGAEA.769615

61. Spreen, G., Kaleschke, L. & Heygster, G. Sea ice remote sensing using AMSR-E 89-GHz channels. Journal of Geophysical Research 113, C02S03 (2008).

62. Cavalieri, D. J. Aircraft active and passive microwave validation of sea ice concentration from the Defense Meteorological Satellite Program special sensor microwave imager. Journal of Geophysical Research 96, 21989–22008 (1991).

63. Stammerjohn, S. E. & Smith, R. C. Opposing Southern Ocean climate patterns as revealed by trends in regional sea ice coverage. Climatic Change 37, 617–639 (1997).

64. Fetterer, F., Knowles, K., Meier, W., Savoie, M. & Windnagel, K. Sea Ice Index, Version 2, updated daily. (2016). doi:http://dx.doi.org/10.7265/N5736NV7

65. Matsuoka, K., Skoglund, A. & Roth, G. Quantarctica [Dataset]. Norwegian Polar Institute (2018). doi:10.21334/npolar.2018.8516e961

66. CCAMLR. European Union proposal to establish the Weddell Sea MPA - CCAMLR-37/29. (2018).

67. CCAMLR. Proposal to establish an East Antarctic Marine Protected Area - CCAMLR-38/21. (2019).

68. Marine Conservation Institute. MPAtlas On-line. (2020). Available at: https://mpatlas.org/.

69. CCAMLR. Delegations of Argentina and Chile. Proposal on a conservation measure establishing a marine protected area in the Domain 1 (Western Antarctic Peninsula and South Scotia Arc). CCAMLR 37-31. (2018).

70. Birdlife International. Aptenodytes forsteri. The IUCN Red List of Threatened Species. The IUCN Red List of Threatened Species 2018 (018).

71. R Core Team. R: A language and environment for statistical computing. R Foundation for Statistical Computing, Vienna, Austria. (2018). Available at: https://www.r-project.org/.

72. . QGIS. (2017). Available at: https://qgis.org/.

