## Supplementary Table 1 for "Juvenile emperor penguin range calls for extended conservation measures in the Southern Ocean"

**Supplementary Table 1. At-sea distribution metrics for the 8 juvenile emperor penguins equipped with ARGOS platforms at Atka Bay.**

| ID | Start trip | End trip | Trip duration (days) | Max distance from colony (km) | Date of max distance from colony | Yearly travelled distance* (km) | Mean daily distance** (km) | %** in ACC area | %** in SO - TL | %** in IUCN - EOO | %** in WSMPA | %** in SGSSI MPA |
|---|---|---|---|---|---|---|---|---|---|---|---|---|
| 65787 | 11/01/19 | 29/12/19 | 352 | 2287 | 21/03/19 | 15593 | 44.3 | 14.4 | 59.3 | 40.1 | 16.9 | 0 |
| 65788 | 13/01/19 | 03/07/19 | 171 | 2390 | 10/04/19 | NA | NA | 28.8 | 44.4 | 36.6 | 10.7 | 5.6 |
| 65789 | 12/01/19 | 24/03/19 | 71 | 1484 | 26/02/19 | NA | NA | 0 | 18.1 | 81.6 | 29.8 | 0 |
| 65790 | 12/01/19 | 12/12/19 | 334 | 2176 | 05/04/19 | 13452 | 40.3 | 16.8 | 54.2 | 42.8 | 5.3 | 0 |
| 65791 | 12/01/19 | 01/06/19 | 140 | 1518 | 20/03/19 | NA | NA | 11.1 | 17.5 | 72.0 | 11.0 | 0 |
| 65792 | 11/01/19 | 13/07/19 | 183 | 1519 | 20/03/19 | NA | NA | 6.5 | 28.8 | 49.8 | 12.7 | 7.5 |
| 65793 | 12/01/19 | 18/06/19 | 157 | 2149 | 04/03/19 | NA | NA | 41.3 | 47.7 | 49.4 | 6.3 | 0 |
| 65794 | 10/01/19 | 01/01/20 | 356 | 2474 | 07/03/19 | 18326 | 51.5 | 19.3 | 48.5 | 50.6 | 5.3 | 0 |
| *Median (date) or Mean values* | *12/01/19* | *08/07/19* | *220.5* | *1999.6* | *20/03/19* | *15790.3* | *45.4* | *18.0* | *55.3* | *48.9* | *10.6* | *1.4* |
| *sd* | *0.9**** | *109.8**** | *110.4* | *421.3* | *15.1**** | *2443.0* | *5.7* | *11.3* | *15.1* | *13.3* | *7.5* | *3.8* |

* Yearly travelled distance was computed using one location every 6 hours (in km), ** % of time spent within the main oceanographic features (ACC, SO-TL) and conservation related areas (IUCN-EOO, MPAs) of the Southern Ocean during the trip duration, *** in days. ACC: Antarctic Circumpolar Current; SO-TL: Southern Ocean Treaty Limits (i.e. at the parallel of 60°S as defined in the Antarctic Treaty); IUCN-EOO: International Union for Conservation of Nature Extent Of Occurrence (i.e. the extent of occurrence of the species considered by the IUCN [33]; WSMPA: Weddell Sea Marine Protected Area; SGSSIMPA: South Georgia and Sandwich Islands Marine Protected Area.
