## Supplementary Fig. 1 for "Juvenile emperor penguin range calls for extended conservation measures in the Southern Ocean"

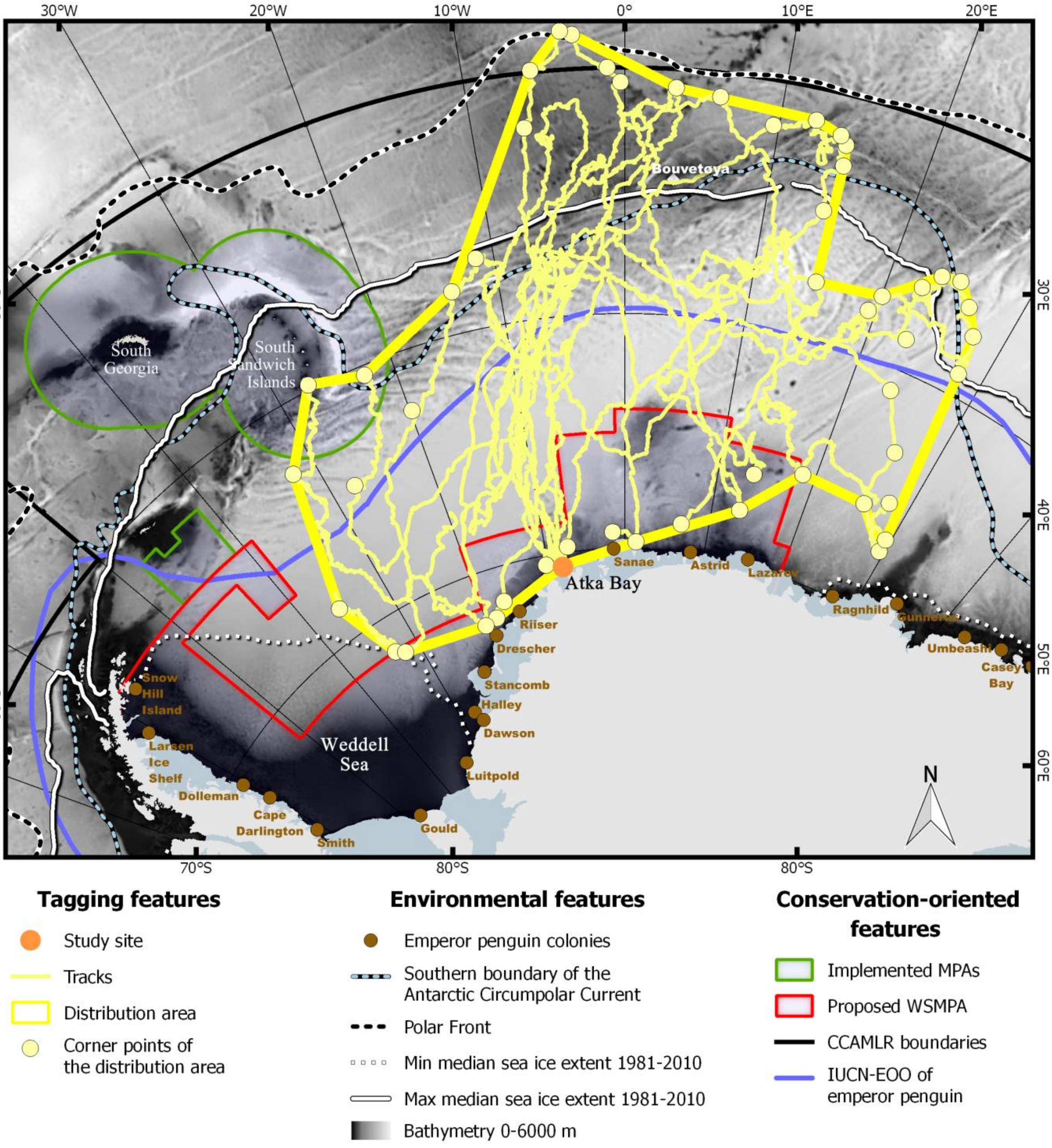


**Supplementary Fig. 1. Distribution over the first year at sea of the 8 juvenile emperor penguins tagged at Atka Bay in 2019**. Main environmental and conservation related features of this sector of the Southern Ocean are specified in the legend. MPAs: Marine Protected Areas; WSMPA: Weddell Sea Marine Protected Area; CCAMLR: Commission for the Conservation of Antarctic Marine Living Resources; IUCN-EOO: International Union for Conservation of Nature Extent Of Occurrence (i.e. the extent of occurrence of the species considered by the IUCN [33].
