## Supplementary Fig. 2 for "Juvenile emperor penguin range calls for extended conservation measures in the Southern Ocean"

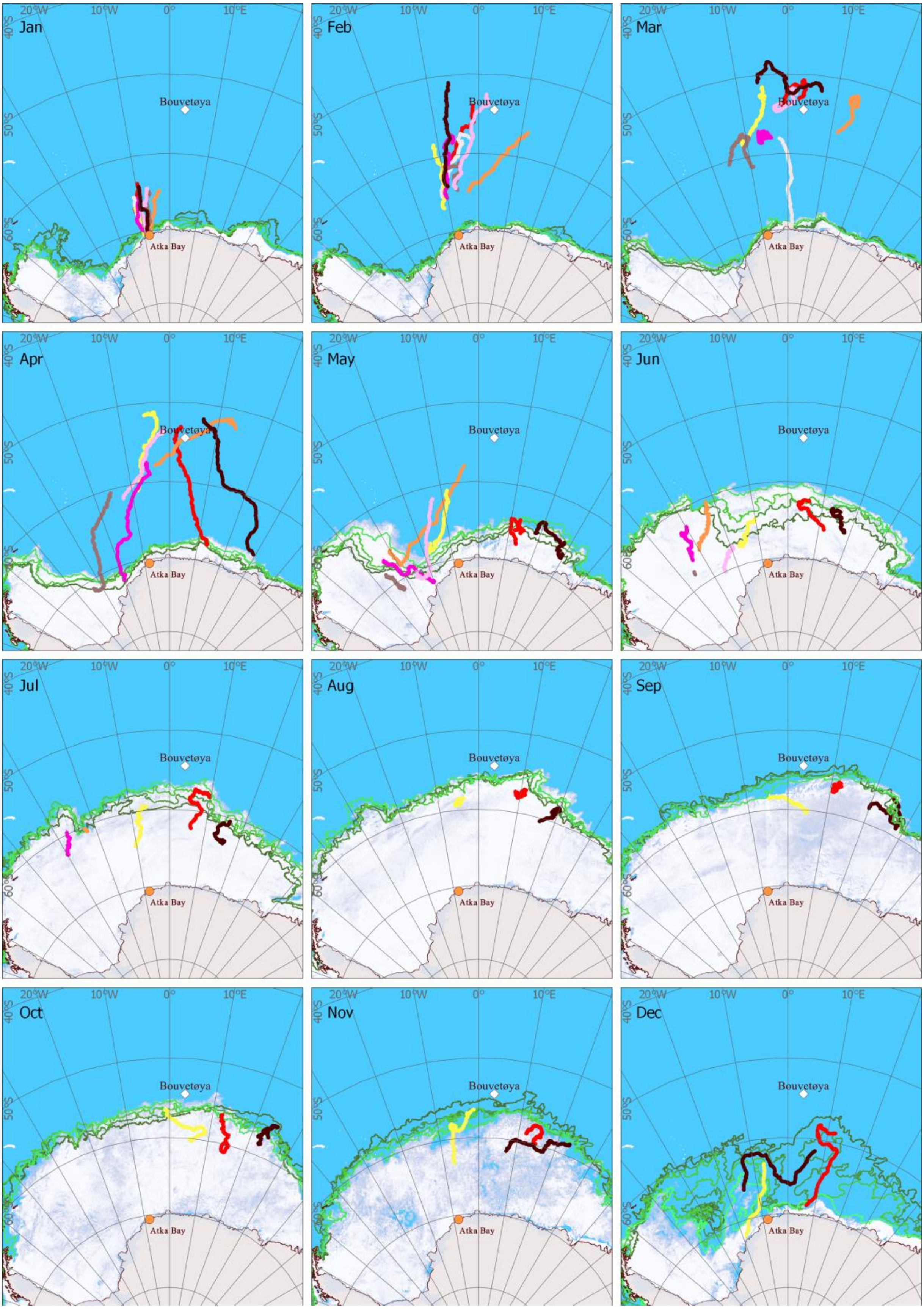


**Supplementary Fig. 2. Monthly tracks of the 8 juvenile emperor penguins tagged at Atka Bay in 2019**. Sea ice concentration is shown for the last day of each month. Sea ice extent is indicated by the green lines for the 4th, 11th, 18th, 25th of each month as indicated by the gradient colour scale from dark green (4th) to light green (25th).
